# The phage shock protein A (PspA) maintains membrane potential and supports NADH dehydrogenase function in mycobacteria

**DOI:** 10.64898/2026.09.09.750162

**Authors:** Pratik Datta, Roberta Provvedi, Charles D. Sohaskey, Rahul Ukey, Julia Puffal, Malavika Prithviraj, Andrea Tellez, Anas Saleh, Rasel Khan, Riccardo Manganelli, Yasu S. Morita, Kyu Rhee, Maria Laura Gennaro

## Abstract

Maintenance of membrane integrity and proton motive force (PMF) is critical for bacterial survival. The phage shock protein (Psp) system, conserved across bacterial species, stabilizes the membrane, maintains PMF, and protects against envelope damage. However, how the conserved effector PspA contributes to PMF maintenance remains unclear. Here, using the mycobacterial Psp system as a genetically tractable model, we provide mechanistic insight into this process. We show that PspA and the accessory protein PspM jointly sustain membrane potential, with PspM required to maintain a ∼70 kDa PspA isoform at the membrane during envelope stress. Loss of PspA increases susceptibility to thioridazine, which targets type II NADH dehydrogenase (NDH-2), and to Ro 48-8071, an inhibitor of menaquinone biosynthesis. Notably, hypersusceptibility to thioridazine is rescued by exogenous menaquinone. Consistent with these phenotypes, a *pspA*-deficient mutant exhibits impaired NADH dehydrogenase activity despite unchanged abundance of NDH-2 and menaquinone (MK-9). Together, these findings identify a functional link between PspA and NADH dehydrogenase-dependent respiration and suggest that PspA contributes to PMF maintenance by supporting respiratory electron transfer.

## Introduction

Bacteria have evolved adaptive responses to diverse stresses, including environmental changes, competition within microenvironments, and host defense mechanisms. A major target of these stresses is the cell envelope, whose integrity is essential for cellular function and survival (1). As the interface with the external environment, the cell envelope regulates the bidirectional movement of substances, controls cell shape and division, and houses stress-sensing systems (1–3). Accordingly, bacteria have evolved dedicated mechanisms to preserve envelope homeostasis under stress. Among these is the phage shock protein (Psp) response, a multigene system induced by envelope stress that stabilizes the cell membrane through coordinated protein interactions (4, 5).

The Psp system, first described as a response to phage infection in *Escherichia coli* (6), centers on the effector protein PspA. PspA is a peripheral membrane protein with a coiled-coil structure that assembles into helical rods (7, 8) and is conserved in eubacteria, archaea, and chloroplasts (5, 9, 10). In *E. coli*, PspA changes its conformation in response to stress, switching from a regulatory oligomer that inhibits its own production to higher-order multimers that exert membrane-stabilizing activity. In both *E. coli* and *Bacillus subtilis*, these transitions are coupled to relocalization of PspA from the cytosol to the inner membrane (4, 11–13). Modulation of PspA conformation and membrane association requires accessory Psp proteins encoded within the same operon (10).

A key feature of the Psp response is its role in maintaining the proton motive force (PMF). Dissipation of PMF has been proposed as a signal for Psp induction (4, 14, 15), and membrane association of PspA is linked to thickening of the lipid bilayer, potentially limiting proton leakage under stress conditions (8, 16, 17). However, the mechanism by which PspA contributes to PMF maintenance remains unclear.

We previously identified a distinct Psp system in mycobacteria and other actinobacteria, which comprises the envelope-stress-responsive transcription factor *clgR* (Rv2745c), the ClgR-regulated *pspA* (Rv2744c), and the gene encoding an integral membrane protein, Rv2743c (renamed as *pspM*) (9, 10, 18). In the *Mycobacterium tuberculosis* complex, the operon also includes a fourth gene, RV2742c (renamed as *pspN*), of unknown function (9, 10, 18). Gene expression dynamics, mutant analyses, and protein–protein interaction data support a model in which interactions between mycobacterial PspA and PspM coordinate the regulatory and membrane-stabilizing functions of PspA (9, 18). These observations suggest that while PspA is the conserved effector, its regulatory and accessory components are phylum-specific and enable system-specific control of PspA activity and membrane targeting (9, 10).

In the present study, we used a defined set of *Mycobacterium smegmatis* mutants spanning the *clgR–psp* operon to dissect the contribution of this system to membrane energetics. We show that the *clgR–pspAM* axis is required for preservation of membrane potential under envelope stress, and define distinct roles for ClgR, PspA, and PspM in this process. We also uncover a functional link between PspA and NADH dehydrogenase activity, indicating that the mycobacterial Psp system contributes to PMF homeostasis, at least in part, by supporting respiratory chain function.

## Results

### PspA and PspM preserve membrane potential under PMF stress

Studies of the Psp system in gram-negative and other bacteria have revealed its role in maintenance of the proton motive force (PMF) (5, 17, 19). Seeking to determine whether this function extends to mycobacteria, we characterized the PMF of isogenic *Mycobacterium smegmatis* strains either containing the native three-gene Psp system (*clgRpspAM*) (20) or bearing in-frame deletion of *clgR* (Δ*clgR*), deletion of both *pspAM* (Δ*pspAM*), or *pspM* (Δ*pspM*) alone (**Fig. S1**). In axenic culture, the Δ*clgR* mutant exhibited a modestly increased doubling time (4.5 h) compared with the wild-type parental strain (3.5 h), whereas the other mutants had doubling times comparable to wild type.

Treatment of exponentially growing *M. smegmatis* wild-type and mutant cultures with carbonyl cyanide m-chlorophenyl hydrazone (CCCP), a protonophore that dissipates PMF (8), at the minimal inhibitory concentration (1x MIC) (**Table S1**) revealed a 2.5- to 4.5-fold reduction in membrane potential in the Δ*clgR*, Δ*pspAM* and Δ*pspM* mutants relative to the isogenic wild-type (WT) strain (**Fig. 1A**), as measured using the membrane potential indicator dye 3,3’-diethyloxacarbocyanine iodide (DiOC_2_) (21). Membrane potential was comparable among untreated WT and mutant strains.

**Fig. 1.**
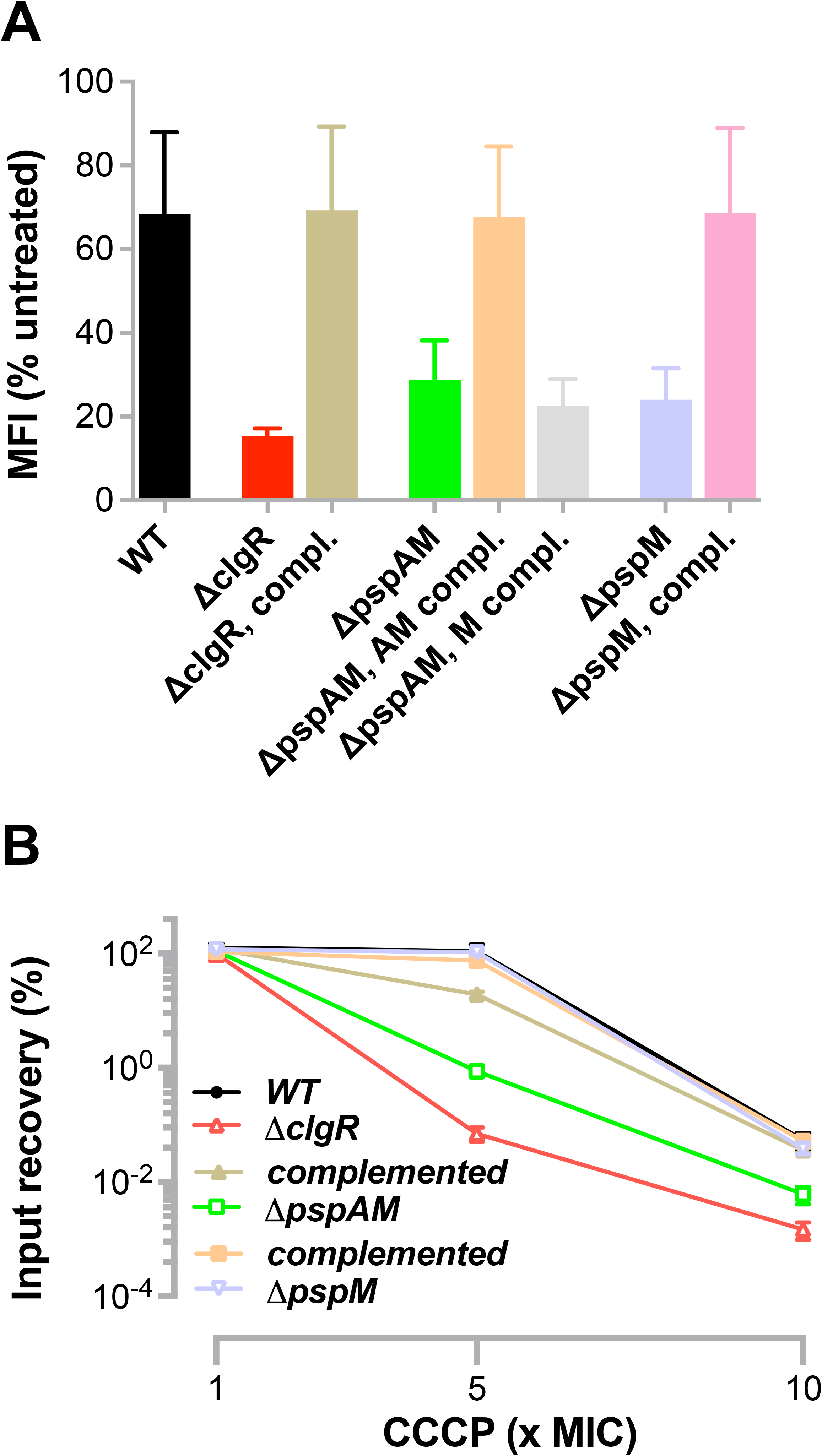
The mycobacterial Psp system protects against CCCP-induced membrane depolarization and loss of viability. **(A)** Membrane potential in *M. smegmatis* wild-type, mutant, and complemented strains following treatment with CCCP (1× MIC) for 3 h, measured by DiOC₂(3) red/green fluorescence ratio. Data are presented as percent change relative to untreated controls. **(B)** Survival of *M. smegmatis* wild-type, mutant, and complemented strains following exposure to increasing concentrations of CCCP for 20 h, determined by CFU enumeration and expressed relative to pretreatment input. In this panel and all subsequent figures, “complemented” always refers to the gene mutation listed above it. Values represent mean ± SEM from three independent experiments.

### Complementation of each mutant with the corresponding

genes restored the WT phenotype following CCCP treatment (**Fig. 1A**). Moreover, expression of *pspM* alone rescued the Δ*pspM* mutant but not the Δ*pspAM* mutant (**Fig. 1A**), indicating that PspA and PspM are jointly required to counter the CCCP-induced membrane depolarization.

Consistent with these results, we found that Δ*clgR* and Δ*pspAM* mutants were significantly more sensitive to CCCP-mediated killing relative to WT cells (>1,500-fold and >100-fold CFU reduction at 5x MIC, respectively) (**Fig. 1B**). The greater susceptibility of the Δ*clgR* mutant relative to the Δ*pspAM* mutant (>10-fold at 5x MIC) suggests that ClgR-regulated genes outside of the *clgR-psp* operon contribute to survival under PMF stress. In contrast, deletion of *pspM* did not affect CCCP lethality (**Fig. 1B**), indicating that membrane depolarization and cell death can be genetically uncoupled.

### PspM supports the membrane association of a distinct PspA species

Since ClgR is the transcription factor that activates the *clgRpsp* operon in mycobacteria (18, 22, 23), the membrane potential defect observed in the Δ*clgR* mutant might be, at least in part, attributable to reduced *pspA* expression. Accordingly, we found that *pspA* mRNA levels in envelope-stressed (CCCP- or SDS- treated) cells were almost two-fold lower in the Δ*clgR* mutant relative to WT cells (**Fig. 2A**). The residual *pspA* induction observed in the Δ*clgR* mutant may be driven by SigE-responsive regulatory sequences upstream of *clgR,* as described in *M. tuberculosis* (22).

**Fig. 2.**
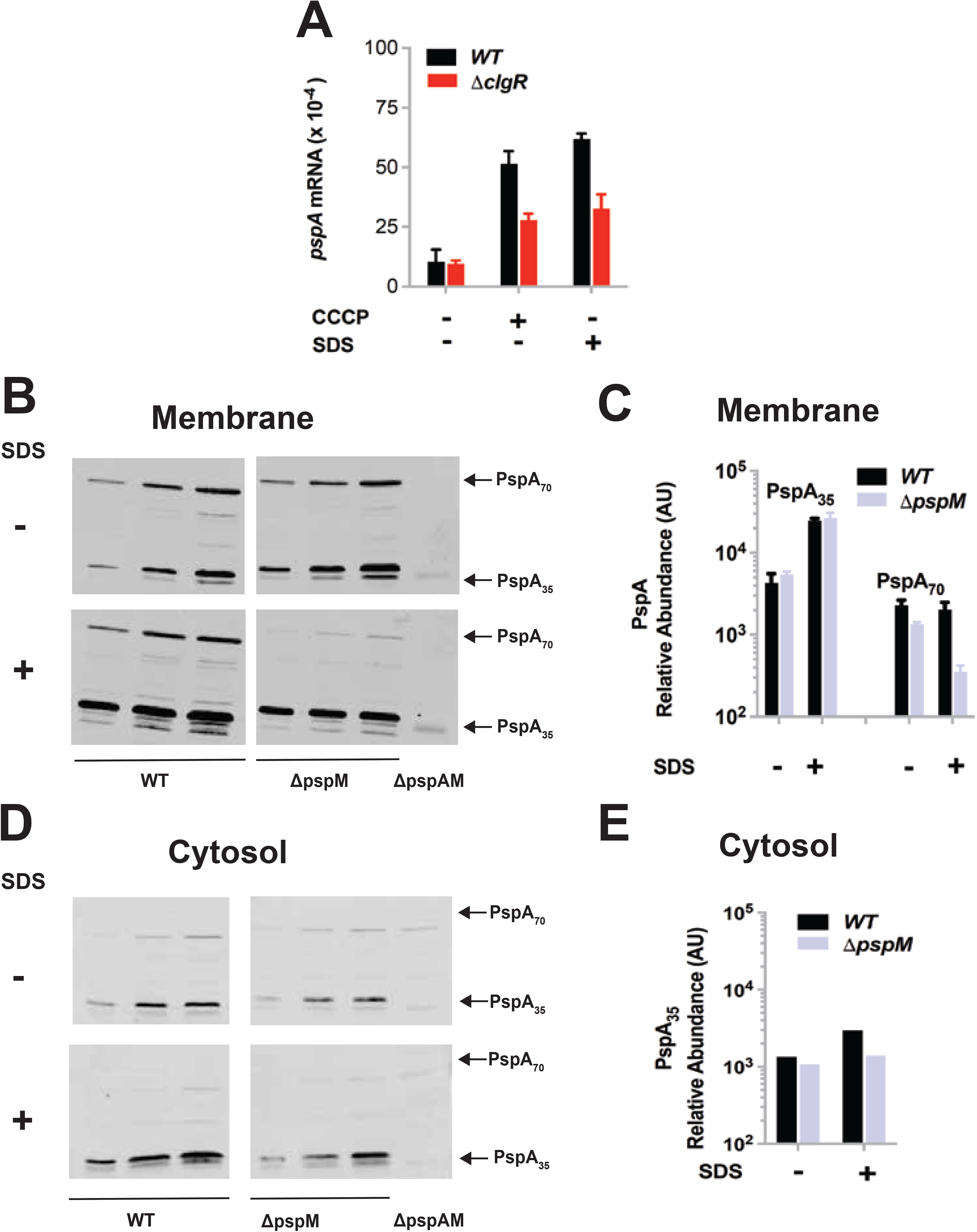
ClgR regulates *pspA* expression and PspM regulates the membrane association of distinct PspA isoforms. **(A)** *pspA* transcript levels in *M. smegmatis* wild-type and Δ*clgR* strains before and after treatment for 45 min with bacteriostatic concentrations of CCCP or SDS. Transcript levels were normalized to 16S rRNA and are presented as mean ± SEM from three independent experiments. **(B)** PspA abundance in membrane fractions of wild-type and Δ*pspM* strains before and after treatment with SDS for 16 h, determined by immunoblotting with anti-PspA antibody. Δ*pspAM* is shown as antibody-specificity control here and in panel D. **(C)** Quantification of membrane-associated PspA isoforms shown in (B), normalized to total protein loading. **(D)** PspA abundance in the corresponding cytosolic fractions of untreated and SDS-treated wild-type and Δ*pspM* strains. The panel also shows the expected position of PspA_70_, which is absent in the cytosol. **(E)** Quantification of cytosolic PspA_35_ shown in (D), normalized to total protein loading. Values in (C) and (E) are mean ± SEM from three lanes per sample.

The impact of CCCP on membrane potential in the Δ*pspM* mutant was, in contrast, more challenging to explain since PspM is an integral membrane protein. We previously reported protein-protein interactions between PspA and PspM in *M. tuberculosis* (18).

This led us to hypothesize that PspM might affect the conformation and/or membrane localization of PspA. To test these possibilities, we used treatment with SDS, an approach often used to study envelope perturbation by the ClgR regulon in *M. tuberculosis* (18, 22). We assessed PspA abundance in untreated and envelope- stressed (SDS-treated) *M. smegmatis* cells by SDS-PAGE and immunoblot analysis of the cell fractions with an anti-PspA hyperimmune serum. By probing the purified membrane-containing fraction, we detected two isoforms of PspA: a ∼35 kDa form (PspA_35_), consistent with the monomer, and a ∼70 kDa form (PspA_70_), likely representing a homo- or hetero-dimeric species (**Fig. 2B**). Upon envelope stress, the levels of PspA_35_ in the membrane fraction were ∼5-fold higher relative to the untreated control; a similar increase in PspA_35_ abundance was observed even in the absence of *pspM* (**Fig. 2BC**). In contrast, levels of the PspA_70_ isoform in the membrane fraction were unaltered by envelope stress but were markedly reduced in the Δ*pspM* mutant (∼4-fold) (**Fig. 2BC**). Moreover, even though PspA remained predominantly membrane-associated, the levels of cytosolic PspA_35_ increased two-fold in response to SDS-mediated stress in WT cells but not in Δ*pspM* mutant cells (**Fig. 2DE**), while the PspA_70_ isoform was undetectable in the cytosolic fraction. Together, these results show that SDS stress induces PspA_35_, leading to membrane-associated PspA_35_ levels that are 10- fold higher than those of PspA_70_. Thus, PspM is not required for the stress-induced increase in membrane-associated PspA35 but is required to maintain PspA70 at the membrane. The moderate decrease in cytosolic PspA_35_ in the Δ*pspM* mutant may reflect redistribution to the membrane during stress.

### PspA sustains NADH dehydrogenase-dependent respiration

Since PMF is primarily maintained by the electron transfer chain (ETC), we next probed the relationship between PspA and ETC function by testing the susceptibility of WT and *clgR-psp* mutants of *M. smegmatis* to chemical inhibitors targeting various components of the respiratory chain [minimal inhibitory concentrations (MIC) for WT and mutant strains are shown in **Table S1**]. Inhibitors of NADH dehydrogenase type I (NDH- 1) (rotenone) and cytochrome bc complex (Q203) were not used because they failed to inhibit growth of *M. smegmatis* (**Table S1**), presumably because the primary NADH dehydrogenase in mycobacteria is NDH-2 rather than NDH-1 (24, 25), and mycobacteria can bypass inhibition of the cytochrome *bc*_1_:aa_3_ complex by transferring electrons from the menaquinone-menaquinol pool directly to the cytochrome *bd-*type menaquinol oxidase (26). We therefore focused on inhibitors of NADH dehydrogenase type II (NDH-2) (thioridazine; THZ), succinate dehydrogenase (SDH) (3-nitrophenol, 3-NP), and ATP synthase (N,N’- dicyclohexylcarbodiimide; DCCD).

When WT and mutant strains were tested with THZ, the Δ*pspAM* mutant exhibited marked hypersensitivity to this drug (>10-fold CFU reduction in the Δ*pspAM* mutant relative to WT at 1.5x MIC) while the Δ*pspM* mutation showed no effect (**Fig. 3A**), indicating that the lethal effect was attributable to loss of *pspA* alone. The moderately increased susceptibility of the Δ*clgR* mutant to THZ (<3-fold CFU reduction in the mutant relative to WT at 1.5x MIC) (**Fig. 3A**) is likely attributable to reduced *pspA* expression in this mutant. When the strain suite was tested with DCCD, hypersensitivity was observed only in the Δ*clgR* mutant but not in the Δ*pspAM* mutant (**Fig. S2A**), pointing to a relationship between ATP synthase function and ClgR-regulated genes outside the Psp system. Moreover, none of the mutants showed altered susceptibility to 3-NP (**Fig. S2B**). Taken together, these data suggest that the <u>ClgR-Psp</u> system is not broadly required for ETC function but is instead specifically linked to NDH-2 activity. NDH-2 catalyzes electron transfer from NADH to menaquinone (MK) (27), the predominant electron carrier to terminal oxidases in mycobacteria (26). We therefore tested the susceptibility of WT and mutant cells to killing by Ro 48-8071, an inhibitor of the menaquinone biosynthetic enzyme MenA (28). Consistent with the THZ results, the Δ*pspAM* mutant, but not Δ*pspM*, exhibited increased susceptibility to Ro 48-8071 (>10- fold CFU reduction Δ*pspAM* relative to WT at 1.5x MIC) (**Fig. 3B**). We next examined whether supplementation with exogenous MK-4 could modify THZ susceptibility. MK-4 suppressed THZ-mediated killing in WT cells and rescued the THZ hypersusceptibility of the Δ*pspAM* mutant in a dose-dependent fashion, reducing its survival defect relative to WT from >8-fold to <3-fold at 0.3 mM MK-4 (**Fig. 3C**). These findings suggested a defect in NADH dehydrogenase activity in *pspA*-deficient cells. Indeed, we found that the Δ*pspAM* mutant, but not the Δ*clgR* or Δ*pspM* mutants, exhibited a 10-fold reduction of NADH dehydrogenase activity and an increased NADH/NAD^+^ ratio relative to WT, the latter being most pronounced following CCCP treatment (∼2-fold) (**Fig. 3DE**). This functional defect was not associated with altered NDH-2 expression or abundance, as both transcript and protein levels were unchanged (**Fig. S3A**), or with reduced steady-state menaquinone abundance, as MK-9, the predominant menaquinone in mycobacteria (29, 30), was detected at similar levels in WT and Δ*pspAM* cells (**Fig. S3B**). Together, these results indicate that loss of PspA affects menaquinone-dependent respiratory function without altering steady-state menaquinone abundance.

**Fig. 3.**
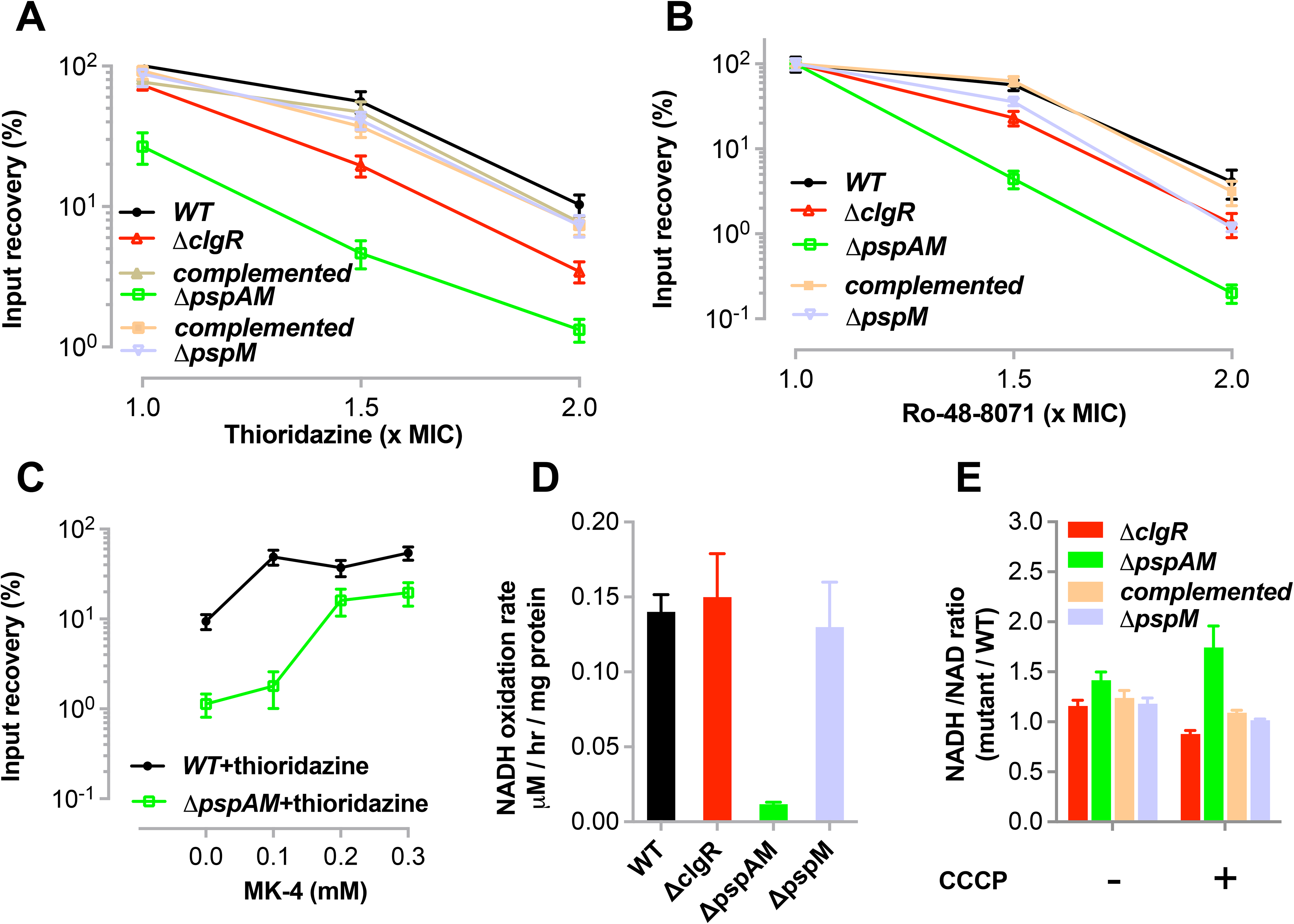
PspA sustains NADH dehydrogenase-dependent respiration. (A) Survival of *M. smegmatis* wild-type, mutant, and complemented strains following 20 h treatment with increasing concentrations of the NDH-2 inhibitor thioridazine. Data are expressed as percent survival relative to pretreatment bacterial counts. **(B)** Survival following 20 h treatment with increasing concentrations of the MenA inhibitor Ro 48-8071. **(C)** Survival of wild-type and Δ*pspAM* strains following treatment with thioridazine (2× MIC) for 20 h in the absence or presence of exogenous menaquinone (MK-4). **(D)** NADH dehydrogenase activity in membrane fractions of wild-type, mutant, and complemented strains, measured as the rate of NADH oxidation and expressed as µmol NADH oxidized per hour per mg of protein. **(E)** NADH/NAD⁺ ratios in wild-type, mutant, and complemented strains following treatment with a bacteriostatic concentration of CCCP for 3 h. Data are expressed as fold change relative to wild type. In all panels, values represent mean ± SEM from three independent experiments.

Protein function can be affected by subcellular localization. We therefore investigated whether the altered phenotype observed in the Δ*pspAM* strain could be explained by NDH-2 mislocalization in the mycobacterial plasma membrane. The mycobacterial plasma membrane is spatially partitioned into functionally distinct domains, termed the inner membrane domain (IMD) and the plasma membrane tightly associated with the cell wall (PM-CW), which can be separated by membrane fractionation by sucrose density gradient centrifugation (31, 32). The IMD is the site of the final steps of MK synthesis, and the proton gradient controls its subcellular localization during membrane perturbation (33, 34). Since NDH-2 was previously detected in the IMD proteome of *M. smegmatis* (32), we investigated whether loss of PspAM altered the localization of NDH-2 within membrane compartments. Fractionation showed that NDH-2 remained enriched in the IMD fraction in the Δ*pspAM* mutant, similar to WT cells (**Fig. S3CD**). These findings indicate that deletion of *pspAM* does not disrupt NDH-2 membrane localization, suggesting that the functional changes observed in the mutant are unlikely to result from mislocalization of the protein within the plasma membrane.

## Discussion

The Psp system has long been known to contribute to PMF maintenance during envelope stress, but the underlying mechanism has remained unclear. Here, we provide mechanistic insight by identifying a functional link between the conserved effector PspA and NADH dehydrogenase activity. Using the mycobacterial system as a model, we show that the *clgR–pspAM* axis is required for maintenance of membrane potential under stress in *Mycobacterium smegmatis*. We define coordinated roles for ClgR, PspA, and PspM and demonstrate that PspA supports NDH-2–dependent respiration. These findings suggest that PspA contributes to PMF homeostasis, at least in part, by sustaining respiratory electron transfer, and raise the possibility that this mechanism may be conserved across Psp systems.

Our data refine the role of PspM by linking it to the functional organization of PspA. In *M. tuberculosis*, PspM modulates ClgR-dependent gene expression, likely through a PspA-ClgR complex (18). Here, we show that PspA exists as a ∼35 kDa monomer and a ∼70 kDa isoform. This result is consistent with a denaturation-resistant homodimer and the propensity of PspA to form higher-order assemblies in other bacterial species (4, 11–13). Whether the PspA₇₀ species represents a homodimer or a higher-order assembly, and whether PspM primarily affects multimerization or its membrane association, remains to be determined. Under basal conditions, PspA is detected in both the cytosol and the membrane fractions, with greater abundance in the latter. Upon envelope stress, membrane-associated PspA₃₅ increases further. PspM is specifically required to maintain the PspA₇₀ isoform at the membrane under envelope stress, and its absence is associated with reduced cytosolic PspA_35_, potentially reflecting altered redistribution of PspA_35_ to the membrane during stress. The distinct effects of *pspM* and *pspAM* deletion -- both impair membrane potential, whereas only loss of PspA affects NADH dehydrogenase-related functions -- suggest that PspM regulates a subset of PspA functions. Together, these findings support a model in which PspM regulates the membrane association of distinct PspA species, thereby modulating PspA function during envelope stress.

A central finding of our work is the link between PspA and NADH dehydrogenase activity. The Δ*pspAM* mutant shows selective hypersensitivity to NDH-2 inhibition, reduced NADH dehydrogenase activity, and altered NADH/NAD⁺ ratios, consistent with impaired respiratory electron transfer. The mutant is also hypersensitive to MenA inhibition, and its hypersensitivity to thioridazine is rescued by exogenous menaquinone. Despite these functional defects, NDH-2 and MK-9 levels are unchanged, suggesting that loss of PspA affects menaquinone-dependent NDH-2 function rather than the abundance of either component. Structural features unique to mycobacterial NDH-2, including a distinct dimer interface and quinone-binding site (35), may render its activity particularly sensitive to membrane organization. In this context, prior evidence linking membrane perturbation to impaired electron flow at the quinone interface (36) raises the possibility that PspA supports NDH-2-menaquinone function by maintaining an appropriate membrane environment. Whether PspA influences NDH-2- menaquinone function through direct interactions with respiratory components or indirectly through effects on membrane organization remains to be determined.

Taken together, our results support a model (**Fig. 4**) in which the mycobacterial Psp system preserves PMF through coordinated mechanisms: ClgR regulates stress-responsive pathways, PspM modulates the membrane association of PspA species, and PspA supports respiratory function. In particular, our findings identify a functional link between PspA and NADH dehydrogenase activity, providing a mechanistic basis for how the Psp system contributes to membrane potential maintenance. This multi-layered organization provides a framework for understanding how stress responses interface with bacterial bioenergetics. While our data support a functional connection between PspA and NDH-2, the molecular basis of this relationship is not defined. Our results also suggest that ClgR contributes to preservation of membrane potential through mechanisms that are not limited to its regulation of the *psp* operon, as the Δ*clgR* mutant exhibits greater sensitivity to CCCP than the Δ*pspAM* mutant. However, the identity of these additional ClgR-regulated factors outside the Psp system remains unknown. Collectively, these findings extend the role of the Psp system in mycobacteria beyond a canonical envelope stress response by linking it to respiratory metabolism. Defining how the Psp system interfaces with respiratory function will provide a mechanistic framework for understanding how mycobacteria maintain bioenergetic homeostasis during membrane stress and may identify vulnerabilities in mycobacterial energy homeostasis with therapeutic potential.

**Fig. 4.**
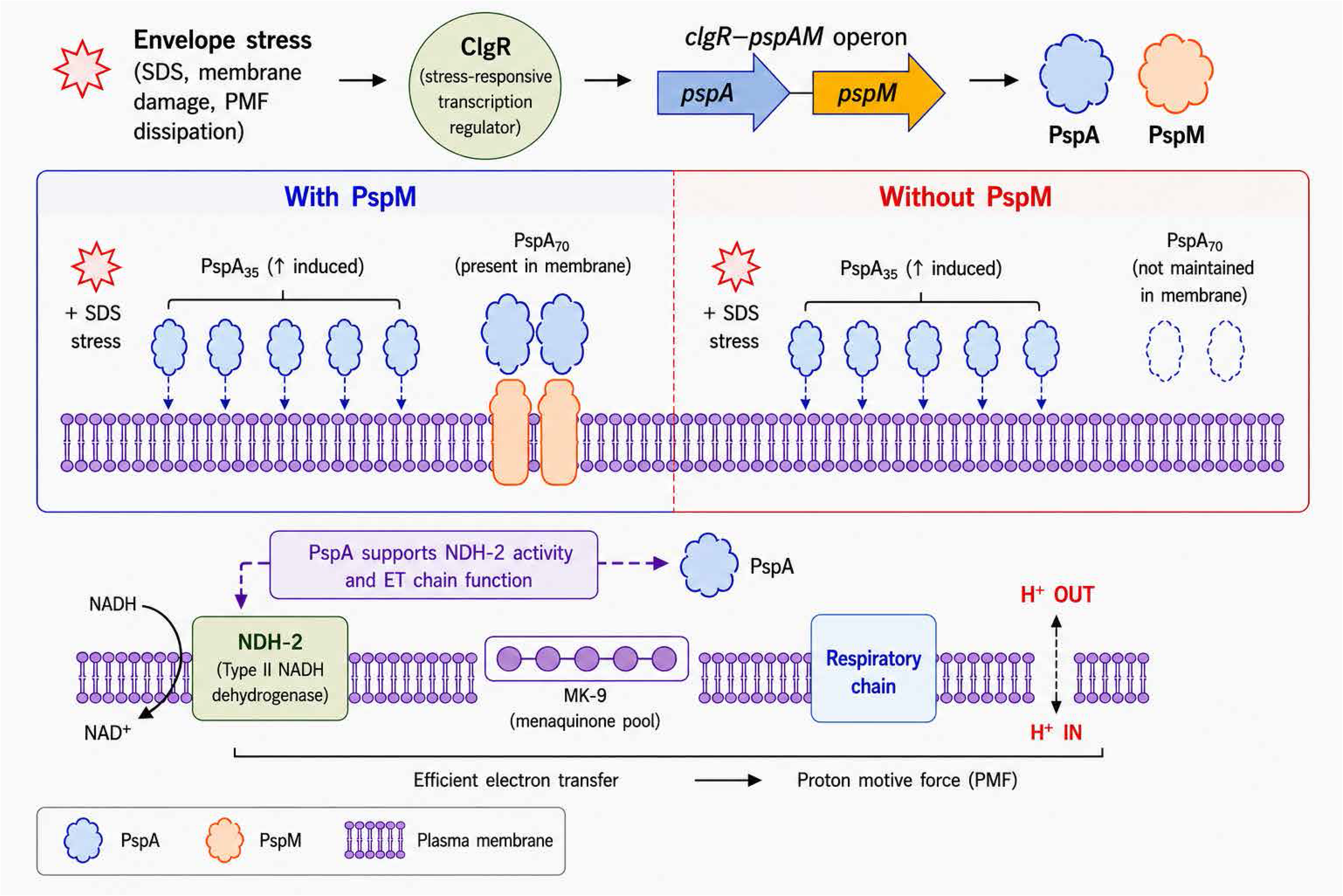
Model for PspA-mediated maintenance of membrane potential through NADH dehydrogenase-dependent respiration. Envelope stress activates the ClgR-regulated *pspAM* operon. PspA₃₅ accumulates at the membrane following SDS stress, whereas maintenance of membrane-associated PspA₇₀ requires PspM. Evidence for a physical interaction between PspA and PspM derives from our previous studies of the *M. tuberculosis clgR–psp* system (18); preferential interaction of PspM with PspA₇₀, as depicted, is inferred from the present findings and remains to be demonstrated. PspA supports NADH dehydrogenase activity and menaquinone-dependent respiratory electron transfer, thereby contributing to maintenance of PMF. The figure was generated using ChatGPT (OpenAI) based on an author-generated draft and subsequently reviewed and edited by the authors.

## Materials and Methods

### Bacterial strains, reagents, and growth conditions

*Escherichia coli* XL1-Blue (Agilent Technologies, Santa Clara, CA) was used for gene cloning. *E. coli* cultures were grown at 37°C in Luria–Bertani (LB) broth or agar (Thermo Fisher Scientific, Waltham, MA), as appropriate. Media were supplemented with kanamycin sulfate (50 μg mL⁻¹) (Thermo Fisher Scientific, Waltham, MA) when required. *M. smegmatis* mc²155 was used to generate knockout mutants and corresponding complemented strains. *M. smegmatis* cultures were grown in 7H9 broth or on 7H10 agar (Difco, Franklin Lakes, NJ). Both solid and liquid media were supplemented with 0.05% Tween 80 and 0.2% glycerol. Liquid cultures were incubated at 37°C with shaking at 200 rpm, while solid plates were incubated at 37°C. For *M. smegmatis* growth, media were supplemented with kanamycin sulfate (Sigma-Aldrich, St. Louis, MO) (25 μg/ml) and hygromycin B (Sigma-Aldrich, St. Louis, MO) (100 μg/ml), as required.

### Construction of *M. smegmatis* knock-out strains

*M. smegmatis* isogenic strains MS110 (Δ*pspAM*) and MS111 (Δ*pspM*) were generated using a recombineering system developed for mycobacteria (37). Briefly, *M. smegmatis* mc^2^155 containing the recombineering plasmid pJV53 was grown overnight at 37°C in Middlebrook 7H9 broth supplemented with 0.2% glycerol, 0.05% Tween-80, 0.2% succinate and 20 µg/ml kanamycin. The following day, cultures were transferred to fresh medium of the same composition and grown to OD_540_ ∼0.4-0.5. After adding acetamide to a final concentration of 0.2%, bacteria were grown for three additional hours and subsequently placed on ice for 1 hour. Electrocompetent cells were prepared as previously described (37).

Targeting linear allelic exchange substrates (AES) were prepared by cloning upstream and downstream homologous regions (∼500 bp) into plasmid pjsc284 [a derivative of pYUB854] (38, 39), flanking the *hyg^R^* cassette. These recombinant plasmids were cleaved with restriction enzymes flanking the homologous regions. After agarose gel purification, 100 ng of AES was electroporated into *M. smegmatis* cells. Transformants were selected on Middlebrook 7H10 plates supplemented with 150 µg/ml hygromycin and the introduced gene deletions were verified by PCR. Primers used for cloning are listed in **Table S2**.

To construct the Δ*clgR* mutant strain (MS133), a 247-bp internal deletion in *MSMEG_2694* (encoding the ClgR transcriptional regulator) was introduced by recombineering in *M. smegmatis* mc²155, as previously described (40). Briefly, two DNA fragments comprising the flanking regions of the target gene were cloned at the borders of an excisable hygromycin (Hyg) resistance cassette flanked by *dif* sites. The resulting construct was introduced by recombineering into an *M. smegmatis* mc²155 derivative carrying pJV53, a replicative plasmid expressing two phage recombinases and conferring kanamycin resistance. Hyg-resistant colonies were selected and verified by PCR. Bacteria were then grown for four generations without selection to allow excision of the Hyg cassette via endogenous Xer recombinases and loss of pJV53. The final unmarked mutant was confirmed by PCR and nucleotide sequencing.

### Construction of complementing plasmids

For complementation of the Δ*clgR* mutant, an 821-bp fragment containing the *clgR* gene and 453 bp of upstream sequence was amplified, flanked with *KpnI* and *HindIII* restriction sites, cloned into the integrative *E. coli–Mycobacterium* shuttle vector pMV306-Kan (41, 42), and introduced into the Δ*clgR* strain. To complement the Δ*pspAM mutant,* a DNA fragment containing *pspAM* and 142 bp upstream of the *clgR co*ding region was amplified from the Δ*clgR* genomic DNA. The resulting construct, which retained the in-frame *clgR* deletion, was cloned into pMV306- Kan between the *KpnI* and *XbaI* sites and electroporated into the Δ*pspAM* strain. For complementation of the Δ*pspM* mutant, two DNA fragments were amplified from Δ*clgR* genomic DNA: (i) a fragment spanning from 142 bp upstream to 258 bp downstream of *clgR*, and (ii) a fragment extending from 47 bp upstream to 16 bp downstream of *pspM*. The second fragment was ligated next to the first fragment to generate a *pspM* complementation construct harboring in-frame deletions of both the *clgR* and *pspA* regions. The final construct was cloned into pMV306-Kan between the *KpnI* and *XbaI* sites and transformed into the Δ*pspM* strain. In all cases, constructs were confirmed by nucleotide sequencing, transformants were selected on appropriate antibiotic-containing media, and correct chromosomal integration was verified by PCR analysis of the vector integration site.

### Minimum inhibitory concentration (MIC) and bactericidal activity of respiratory chain inhibitors

Exponentially growing *M. smegmatis* cultures were adjusted to an OD₆₀₀ of 0.01 (approximately 1 × 10 colony-forming units (CFU) per ml) and inoculated into 96-well microtiter plates containing two independent series of two-fold dilutions of the indicated compounds in 7H9 broth. Wells lacking compounds served as growth controls. Plates were incubated at 37°C with shaking for 24 h, after which bacterial growth was evaluated by measuring turbidity at OD₆₀₀. The minimum inhibitory concentration (MIC) was defined as the lowest concentration that inhibited detectable growth relative to untreated control.

Bactericidal activity was assessed by determining bacterial survival following exposure to the indicated multiples of the MIC. Mid-log-phase cultures of *M. smegmatis* wild-type, mutant, and complemented strains were adjusted to approximately 1 × 10 CFU/ml and exposed to the indicated compounds at 37°C for 16 h with shaking. Following treatment, bacterial survival was quantified by CFU enumeration and compared with the corresponding pretreatment inoculum.

### Membrane depolarization assay

Exponentially growing cultures of *M. smegmatis* wild-type and mutant strains were either left untreated or treated for 3 h with CCCP at 1× MIC concentration (**Table S1**). Following treatment, cultures were incubated at 37°C without shaking. During the final 30 min of incubation, the membrane potential–sensitive carbocyanine dye DiOC₂(3) (Molecular Probes) was added to both treated and untreated cultures. The red-to-green fluorescence ratio of DiOC₂(3), which reflects membrane potential, decreases upon CCCP-mediated membrane depolarization (43). Mean fluorescence intensity (MFI) in the red and green channels was measured by flow cytometry using a BD LSR Fortessa X-20 (BD Biosciences, San Jose, CA), and red-to-green fluorescence ratios were calculated. Percent changes in fluorescence ratios for CCCP-treated cultures were determined relative to untreated controls. All experiments were performed using three biological replicates.

### Envelope stress treatments (CCCP and SDS

Bacteriostatic conditions were defined as the highest concentration of SDS or CCCP that did not reduce the viability of wild-type or mutant strains after 16 h of exposure, relative to pretreatment samples collected at OD_600_=0.3. Based on these assays, 0.03% SDS and 15 μM CCCP were selected as bacteriostatic concentrations.

For *pspA* gene expression analysis, *M. smegmatis* cultures were treated with the indicated bacteriostatic concentrations for 45 min, after which 1 mL aliquots were collected for RNA extraction. For PspA protein analysis, cultures were treated with 0.03% SDS for 16 h. Bacterial pellets collected before and after treatment were lysed and fractionated for downstream protein analysis.

### Transcript enumeration

Exponentially growing *M. smegmatis* WT and mutant cultures were treated with bacteriostatic concentrations of CCCP or SDS, as mentioned above. Bacterial pellets were resuspended in 1 mL TRI Reagent (Molecular Research Center, Cincinnati, OH), and cells were disrupted with 0.1 mm zirconia beads (BioSpec Products, Inc., Bartlesville, OK) using a Mini-Beadbeater-16 (BioSpec Products, Inc.). RNA extraction was performed as previously described (18). Complementary DNA (cDNA) synthesis was carried out using ThermoScript Reverse Transcriptase (Thermo Fisher Scientific, Waltham, MA). Quantitative real-time PCR (qRT-PCR) was performed using AmpliTaq Gold DNA Polymerase (Applied Biosystems, Waltham, MA ) in a Stratagene Mx4000 (Agilent Technologies, Santa Clara, CA). Transcript levels were quantified using gene-specific molecular beacons and normalized to the *M. smegmatis* 16S rRNA transcript levels, as previously described (44).

### Subcellular fractionation

Exponentially growing *M. smegmatis* cultures were treated with 0.03% SDS for 16 h. Pre- and post-treatment pellets were resuspended in PBS containing protease inhibitors and lysed with 0.1 mm zirconia beads using 10 cycles of bead beating (1 min each) with intermittent cooling on ice. Lysates were clarified by sequential centrifugation at 10,000 rpm and 15,000 rpm using a refrigerated Eppendorf benchtop centrifuge (Hamburg, Germany). The resulting supernatants were then subjected to ultracentrifugation (Beckman Coulter, Brea, CA) at 100,000 × g for 90 min to separate the membrane fraction from the cytosolic fraction. Membrane pellets were washed with PBS, solubilized overnight at 4°C in B-PER (Thermo Fisher Scientific, Waltham, MA), and protein concentrations were determined by BCA assay prior to SDS-PAGE and immunoblotting.

Proteins were transferred to PVDF membranes and probed with rabbit anti-PspA antibody; a separate immunoblot for NDH-2 was performed on the membrane fraction of mid-log cells using mouse anti-His antibody (Thermo Fisher Scientific, Waltham, MA). Immunoblots were probed with the appropriate secondary antibodies. Signals were detected using the LI-COR Odyssey system (LI-COR Biosciences, Lincoln, NE) and quantified with Odyssey software v1.2. Band intensities were normalized to total protein per lane, measured using REVERT Total Protein Stain (LI-COR Biosciences, Lincoln, NE).

### Separation of membrane domains and immunoblotting

Subcellular fractions were obtained by sucrose gradient fractionation of cell lysates as previously described (31, 32). For immunoblotting, equal volumes of subcellular fractions were separated by SDS-PAGE, transferred to a PVDF membrane, and analyzed for NHD-2 abundance using mouse anti-His antibodies. PimB’ and MptA were used as IMD and PM-CW markers, respectively, and the corresponding antibodies were raised in rabbits against PimB’ and MptA peptides, as previously described (45).

### NAD+ and NADH measurements

*M. smegmatis* wild-type, mutant, and complemented strains were grown to OD₆₀₀ = 0.5-0.7 and treated with a bacteriostatic concentration of CCCP for 3 h. Untreated and treated samples were collected for NAD⁺ and NADH measurements as described previously (46). Briefly, cell pellets were extracted in 0.2 M HCl (NAD⁺) or 0.2 M NaOH (NADH) by incubation at 55°C for 10 min, followed by neutralization with 0.1 M NaOH or 0.1 M HCl, respectively. Extracts were clarified by high-speed centrifugation. NAD⁺ and NADH levels were quantified using an enzymatic cycling assay in 96-well plates. A master mix for the cycling cocktail was prepared by mixing 1 ml each of 1 M bicine (pH 8.0), 100% ethanol, 40 mM EDTA (pH 8.0), and 4.2 mM thiazolyl blue tetrazolium bromide (MTT), and 2 ml of 16.6 mM phenazine ethosulfate (PES). A 50 μl aliquot of the cocktail was mixed with 45 μl of sample. The mixture was briefly incubated at 30^0^C, followed by addition of 5 μl of alcohol dehydrogenase (500 U/ml in 0.1 M bicine pH 8.0). Absorbance at 550 nm was recorded over 20 min at 2-min intervals. Concentrations of NAD^+^ or NADH were measured by following the reduction of MTT by the dehydrogenase in the presence of PES. Concentrations were calculated from standard curves included in the same plate. NADH/NAD⁺ ratios were determined from three biological replicates, and fold changes relative to the wild-type strain were calculated.

### NADH dehydrogenase activity

Mid-log-phase *M. smegmatis* cultures were harvested, washed, and resuspended in phosphate buffer (50 mM K_2_HPO_4_ pH 7.5; 5 mM MgSO_4_). A DNase and proteinase inhibitor cocktail was added to the bacterial suspension prior to lysis by French press (1,000 lb/in^2^/cm^2^). Subcellular fractions were separated by differential centrifugation, and purified membrane fractions were subsequently obtained by ultracentrifugation as described above. Fragmented membrane fractions were resuspended in phosphate buffer, and protein concentrations were determined as mentioned above. NADH dehydrogenase activity was assayed by measuring the rate of NADH oxidation as the decrease in absorbance at 340 nm, using a fixed amount of exogenously added NADH and 2 mM menadione (type II), and a Libra S32PC spectrophotometer at room temperature, as previously described (47). A control reaction with heat-inactivated membrane extracts was included. Activities were determined from the linear section of the curve, and specific activity was expressed as µmol of NADH oxidized per hour per mg of protein.

### Menaquinone (MK-9) quantification

*M. smegmatis* strains were grown in 7H9 broth supplemented with 0.02% glycerol and 0.05% Tween 80 to OD=1, from which 1 ml was seeded onto 0.2 micron pore size nylon filters (Millipore) and placed atop 7H10 agar media. Bacteria laden filters were incubated for an additional 2 days at 37^0^C, after which they were plunged into glass tubes containing 7 ml of a 2:1 chloroform:methanol mixture and allowed to incubate for 24 hours with gentle shaking at 100 rpm at 37^0^C. Samples were subsequently centrifuged at 4,000 rpm for 15 min at 4^0^C, after which the supernatant was collected and evaporated to dryness using a GeneVac EZ-2 evaporator. The residual material was rinsed with 2 mL of chloroform:methanol (1:1, v/v), followed by sonication and vortexing, and subsequently centrifuged at 4,000 rpm for 15 min at 4 °C. The resulting supernatant was collected and transferred to a pre- weighed 4 mL amber vial. Lipid extracts were dried under vacuum and resuspended to a final concentration of 1 mg mL⁻¹ in chloroform:methanol (1:1, v/v).

For LC–MS analysis, 150 µL of the 1 mg mL⁻¹ lipid stock was dried under vacuum and resuspended in 150 µL of hexane:isopropanol (70:30, v/v) containing 0.02% (m/v) formic acid and 0.01% (m/v) ammonium hydroxide. Samples were centrifuged at 4,000 rpm for 15 min, and 100 µL of the supernatant was transferred to an amber glass LC vial fitted with a glass insert and sealed with a cap.

Lipid analysis was performed by liquid chromatography–mass spectrometry (LC–MS) as described by Layre et al (48). Samples (10 µL injection volume) were separated on an Agilent 1260 Infinity II LC system using an Inertsil Diol column (2.1 × 150 mm, 3 µm; GL Sciences). Mobile phase A consisted of hexane:isopropanol (70:30, v/v) containing 0.02% (v/v) formic acid and 0.04% (v/v) ammonium hydroxide, and mobile phase B consisted of isopropanol:methanol (70:30, v/v) containing 0.02% (v/v) formic acid and 0.04% (v/v) ammonium hydroxide. The flow rate was maintained at 0.150 mL min⁻¹. The 44-min normal-phase gradient was as follows: 0–10 min, 0% B; 17–22 min, 50% B; 30– 35 min, 100% B; and 40–44 min, 0% B, followed by a 6-min post-run equilibration at 0% B. Mass spectrometric data were acquired on an Agilent 6545 Q-TOF LC/MS (Agilent Technologies) equipped with an Agilent Jet Stream electrospray ionization source, operated in extended dynamic range in both positive and negative ion modes. The capillary voltage was set to 3,500 V and the nozzle voltage to 1,000 V. The nebulizer pressure was 30 psi, with a sheath gas flow of 11 L min⁻¹ at 350 °C. The drying gas was maintained at 325 °C with a flow rate of 5 L min⁻¹. The fragmentor, skimmer, and octopole radio frequency voltages were set to 200 V, 65 V, and 650 V (Vpp), respectively. Data were acquired in centroid mode at 1 spectrum s⁻¹ over an m/z range of 50–3,000. Analysis of MK-9 levels was determined by integrating the total area under the curve corresponding to targeted extracted ion chromatograms of the accurate mass-retention time identifier for the [M+H] and [M+NH4] adducts of MK-9 reported in the MycoMap (48).

## Acknowledgements

We thank Serena Guggia and Fedora Babic for technical support in the construction of the *M. smegmatis* mutants and Karl Drlica for critical comments on the manuscript. This work was funded in part by NIH R01HL149450 and R21AI153660 (MLG); NIH R21AI144748 and a grant from Pittsfield Anti-Tuberculosis Association (YSM); NIH R01AI140472 (KR); and fellowships from the University of Massachusetts Amherst as part of the Chemistry-Biology Interface Training Program (National Research Service Award T32 GM008515 and GM139789) (MP) and Science Without Borders from CAPES-Brazil (0328-13-8) (JP).

## Supplementary Figure legends

**Fig. S1. Schematic representation of the *clgR–pspAM* locus in wild-type, mutant, and complemented strains.** The Δ*clgR* mutant contains an unmarked in-frame deletion, whereas the *pspAM* and *pspM* regions were replaced with a hygromycin resistance cassette (*hygR*). “CompL” denotes the corresponding chromosomally integrated complementation constructs. The Δ*pspAM* mutant was complemented with either *pspAM* or *pspM* alone.

**Fig. S2. ClgR, but not the Psp system, protects against ATP synthase inhibition. (A)** Survival of *M. smegmatis* wild-type and mutant strains following 20-h exposure to increasing concentrations of the ATP synthase inhibitor DCCD. **(B)** Survival following 20-h exposure to increasing concentrations of the succinate dehydrogenase inhibitor 3- nitrophenol (3-NP) for 20 h. Data are expressed as percent survival relative to pretreatment bacterial counts. Values represent mean ± SEM from three independent experiments.

**Fig. S3. Loss of PspA does not alter NDH-2 abundance or membrane localization or steady-state MK-9 levels. (A)** *ndh-2* transcript levels (left) and membrane-associated NDH-2 protein abundance (right) in wild-type and Δ*pspAM* strains. Transcript levels were normalized to 16S rRNA. NDH-2 abundance was normalized to total protein. **(B)** Steady-state MK-9 levels in wild-type and Δ*pspAM* strains. Relative MK-9 abundance was determined as described in Methods and expressed as the integrated area under the curve for the [M+H] and [M+NH₄] adducts of MK-9. Values represent mean ± SEM of three biological replicates per strain. **(C, D)** Distribution of NDH-2 between the inner membrane domain (IMD) and plasma membrane–cell wall (PM-CW) fractions of wild-type and Δ*pspAM* strains. PimB′ and MptA served as markers for the IMD and PM-CW fractions, respectively. For immunoblot analyses in this figure, NDH-2 was expressed as a His-tagged protein (NDH-2–6×His) and detected using an anti-His antibody.

