## Supplementary Information for "The phage shock protein A (PspA) maintains membrane potential and supports NADH dehydrogenase function in mycobacteria"

#### Supplementary Figure legends

**Fig. S1. Schematic representation of the *clgR–pspAM* locus in wild-type, mutant, and complemented strains.** The Δ*clgR* mutant contains an unmarked in-frame deletion, whereas the *pspAM* and *pspM* regions were replaced with a hygromycin resistance cassette (*hygR*). "CompL" denotes the corresponding chromosomally integrated complementation constructs. The Δ*pspAM* mutant was complemented with either *pspAM* or *pspM* alone.

**Fig. S2. ClgR, but not the Psp system, protects against ATP synthase inhibition. (A)** Survival of *M. smegmatis* wild-type and mutant strains following 20-h exposure to increasing concentrations of the ATP synthase inhibitor DCCD. **(B)** Survival following 20-h exposure to increasing concentrations of the succinate dehydrogenase inhibitor 3-nitrophenol (3-NP) for 20 h. Data are expressed as percent survival relative to pretreatment bacterial counts. Values represent mean ± SEM from three independent experiments.

**Fig. S3. Loss of PspA does not alter NDH-2 abundance or membrane localization or steady-state MK-9 levels. (A)** *ndh-2* transcript levels (left) and membrane-associated NDH-2 protein abundance (right) in wild-type and Δ*pspAM* strains.

Transcript levels were normalized to 16S rRNA. NDH-2 abundance was normalized to total protein. **(B)** Steady-state MK-9 levels in wild-type and Δ*pspAM M. smegmatis* strains. The relative abundance of MK-9 was determine as described in Methods and reported as integrated area under the curve values corresponding to the [M+H] and [M+NH4] adducts of MK-9. Values represent the average of the biological replicates for each strain. **(C, D)** Distribution of NDH-2 between the inner membrane domain (IMD) and plasma membrane–cell wall (PM-CW) fractions of wild-type and Δ*pspAM* strains. PimB′ and MptA served as markers for the IMD and PM-CW fractions, respectively. For immunoblot analyses in this figure, NDH-2 was expressed as a His-tagged protein (NDH-2–6×His) and detected using an anti-His antibody.

#### Supplementary Tables

**Supplementary Table 1.** Minimum inhibitory concentrations of compounds used in this study against wild-type and mutant *M. smegmatis*.

**Supplementary Table 2.** List of primers for mutant construction.

### Figure S1

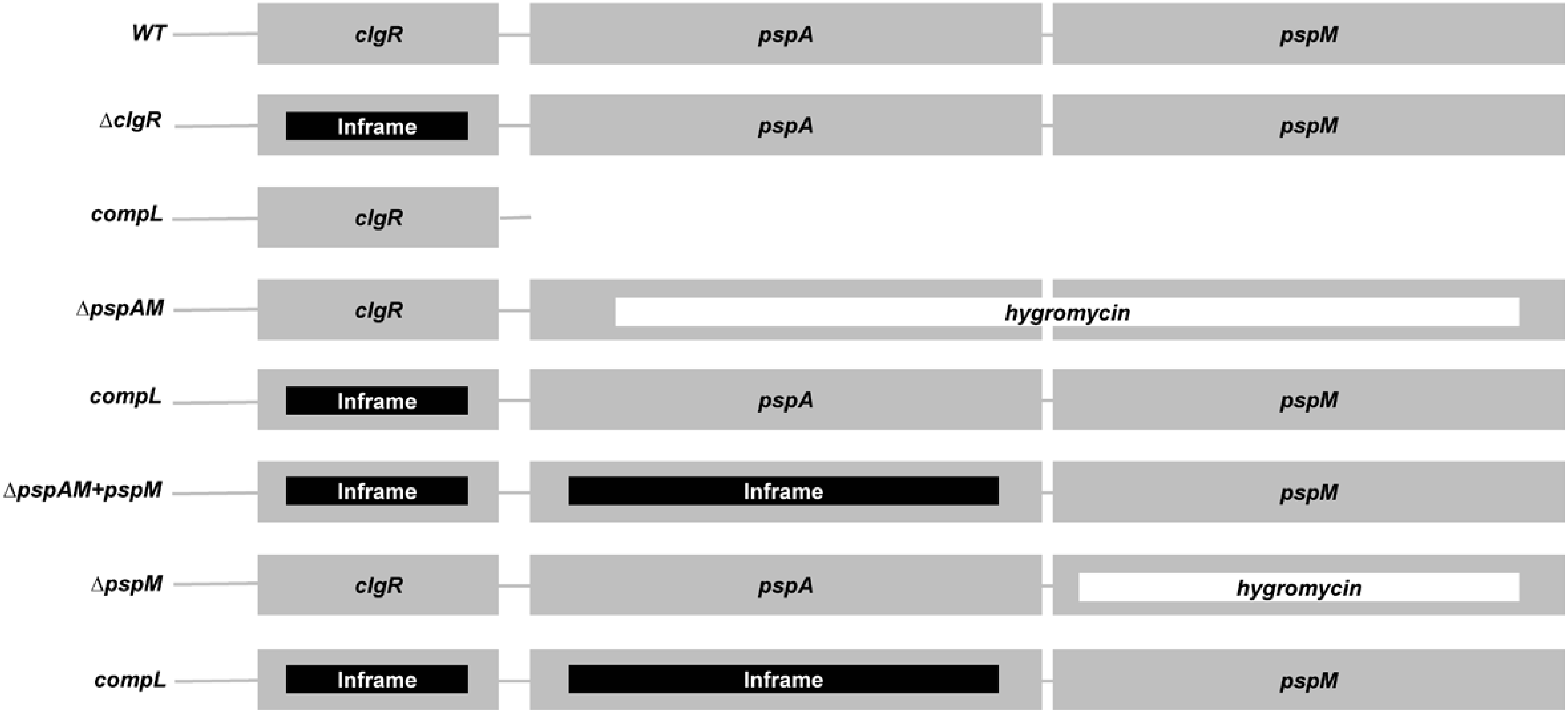

WT
clgR
pspA
pspM
ΔclgR
Inframe
pspA
pspM
compL
clgR
ΔpspAM
clgR
hygromycin
compL
Inframe
pspA
pspM
ΔpspAM+pspM
Inframe
Inframe
pspM
ΔpspM
clgR
pspA
hygromycin
compL
Inframe
Inframe
pspM

### Figure S2

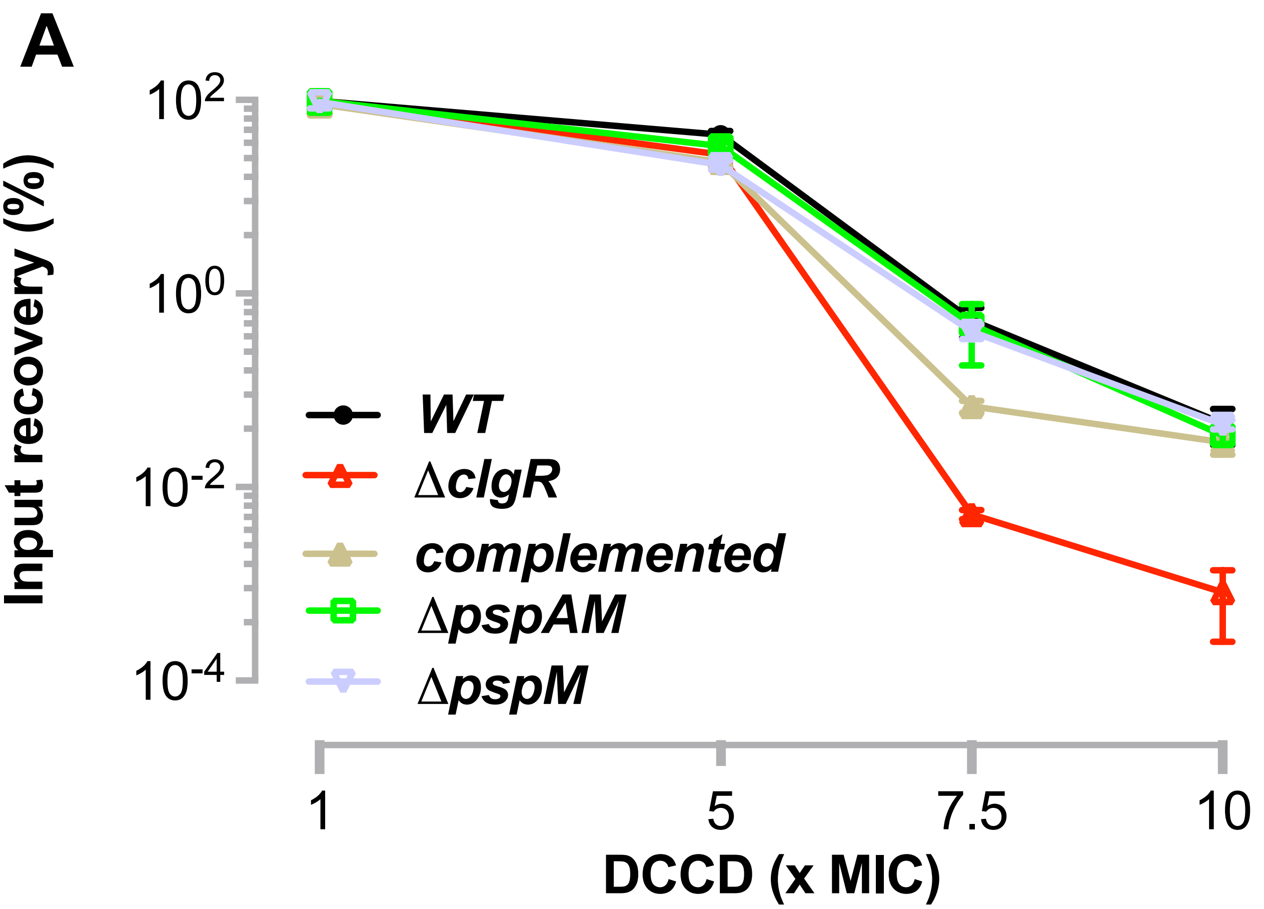

A
Input recovery (%)
$10^2$
$10^0$
$10^{-2}$
$10^{-4}$
WT
ΔclgR
complemented
ΔpspAM
ΔpspM
1
5
7.5
10
DCCD (x MIC)


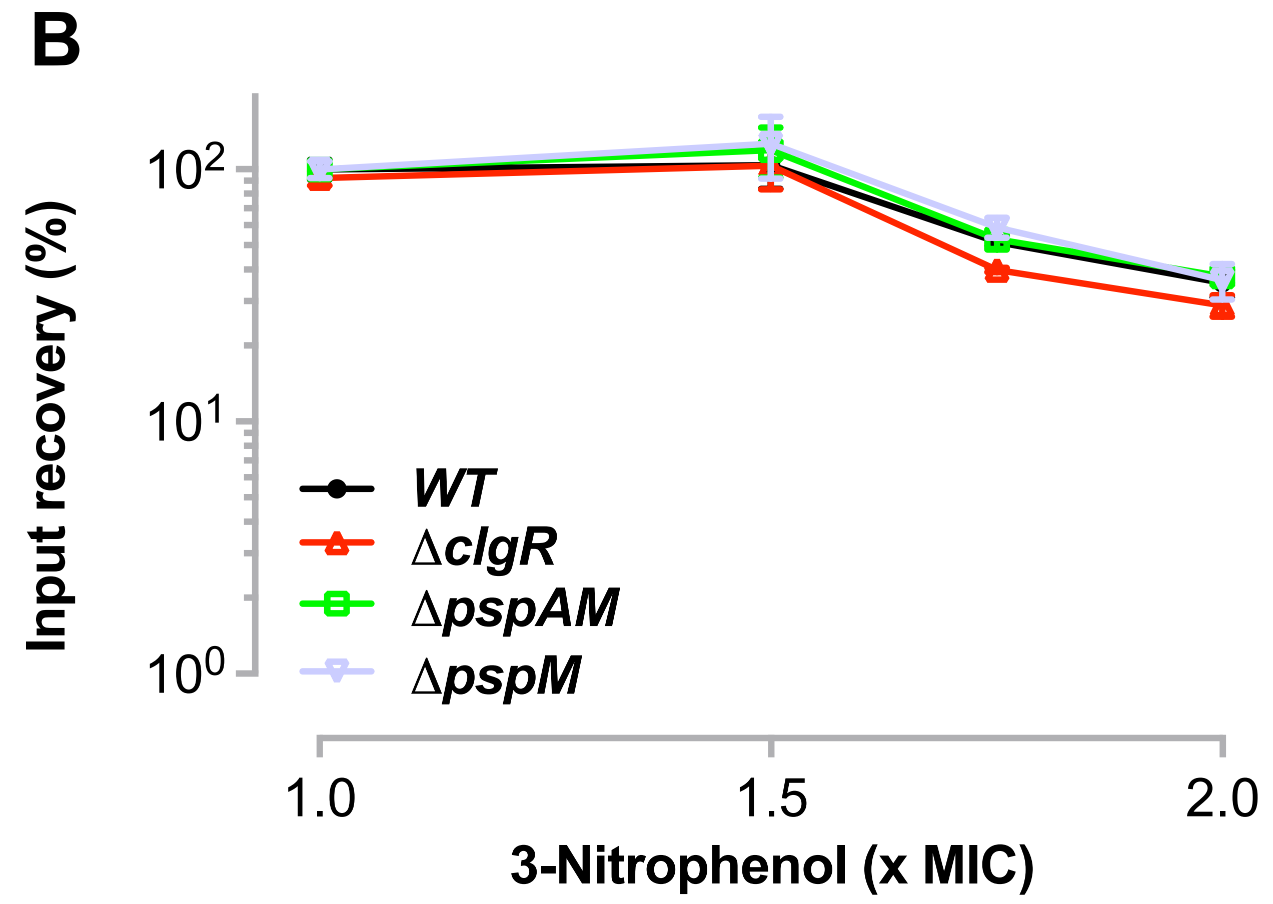

B
Input recovery (%)
$10^2$
$10^1$
$10^0$
WT
ΔclgR
ΔpspAM
ΔpspM
1.0
1.5
2.0
3-Nitrophenol (x MIC)

### Figure S3

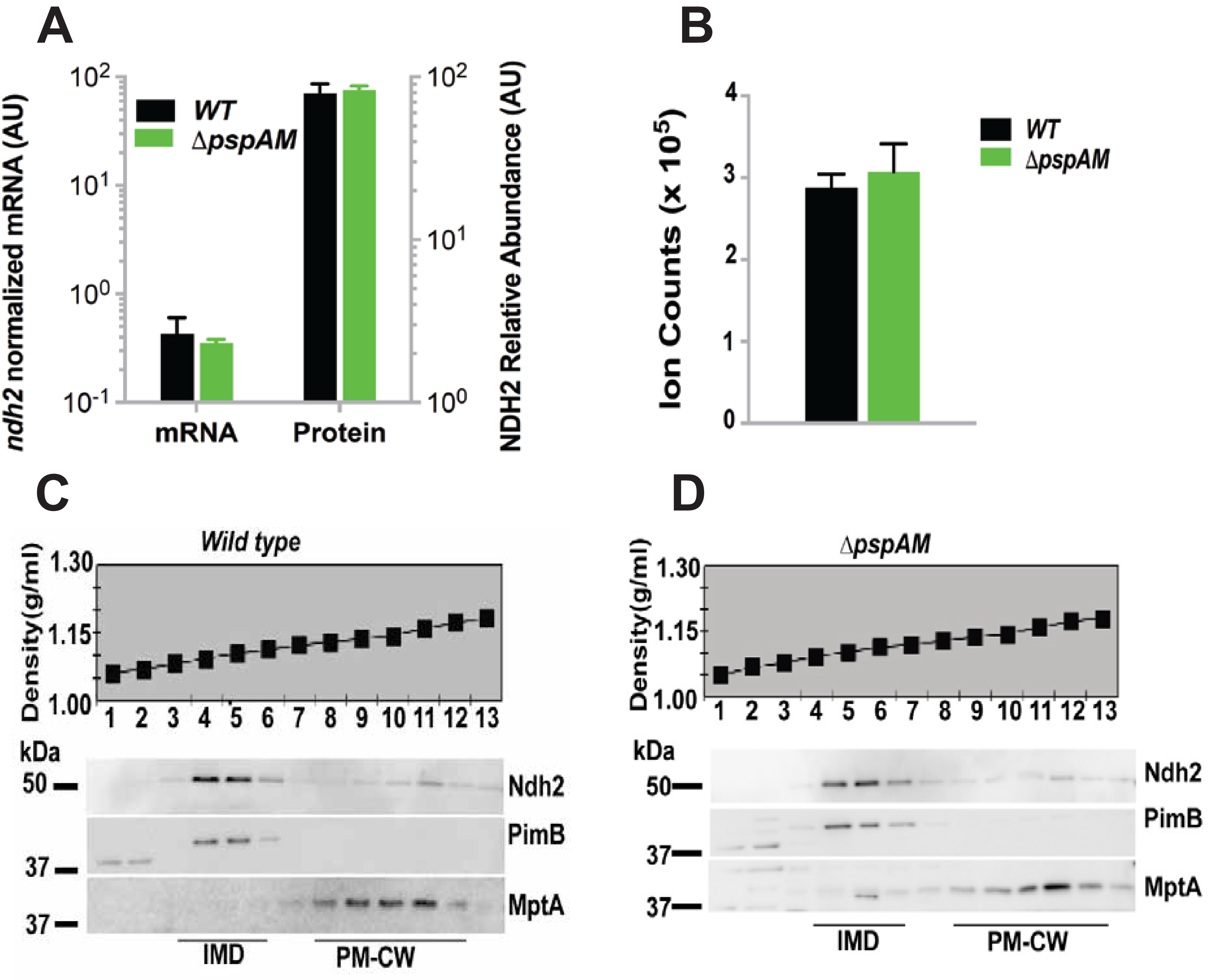

A
B
C
D
WT
ΔpspAM
ndh2 normalized mRNA (AU)
NDH2 Relative Abundance (AU)
mRNA
Protein
Ion Counts (x 10^5)
Wild type
Density(g/ml)
kDa
Ndh2
PimB
MptA
IMD
PM-CW

**Supplementary Table 1. Minimum inhibitory concentrations of compounds used in this study against wild-type and mutant *M. smegmatis*.**

| **Inhibitors** | **Mc$^2$155** | **Δ*pspAM*** | **Δ*clgR*** |
|---|---|---|---|
| CCCP (μM) | 3 | 3 | 2.5 |
| Thioridazine (μg/ml) | 10 | 7 | 10 |
| DCCD (μg/ml) | 4 | 4 | 4 |
| Q203 (μg/ml) | >16 | >16 | ≥16 |
| 3-Nitrophenol (μg/ml) | 128 | 128 | 128 |
| Rotenone (μg/ml) | >256 | >256 | >256 |

Minimum inhibitory concentrations of wild-type (Mc$^2$155) and mutant *M. smegmatis* were determined as described in Materials and Methods. Lack of inhibition by rotenone, an inhibitor of the type I NADH dehydrogenase (NDH-1), is consistent with reliance of mycobacteria on rotenone-insensitive NDH-2 for NADH oxidation (1). Lack of inhibition by Q203, an inhibitor of the cytochrome *bc1* complex, reflects respiratory flexibility, whereby electron flux can be rerouted through Q203-insensitive cytochrome *bd* oxidase (2).

1. Weinstein EA, Yano T, Li LS, Avarbock D, Avarbock A, Helm D, McColm AA, Duncan K, Lonsdale JT, Rubin H. 2005. Inhibitors of type II NADH:menaquinone oxidoreductase represent a class of antitubercular drugs. Proc Natl Acad Sci U S A.
2. Kalia NP, Hasenoehrl EJ, Ab Rahman NB, Koh VH, Ang MLT, Sajorda DR, Hards K, Gruber G, Alonso S, Cook GM, Berney M, Pethe K. 2017. Exploiting the synthetic lethality between terminal respiratory oxidases to kill Mycobacterium tuberculosis and clear host infection. Proc Natl Acad Sci U S A 114:7426-7431.

**Supplementary Table 2. List of primers for mutant construction**.

| Amplified region | Upper primer sequence | Lower primer sequence | Fragment length |
|---|---|---|---|
| Upstream *ΔpspAΔMSMEG_2696* (MS110) | <u>GGTACC</u>GAACCGATAAGTTGGCACCACAA | <u>TCTAGA</u>GTTCTGCTCGACGGCCTTCTT | 509 bp |
| Downstream *ΔpspAΔMSMEG_2696* (MS110) | <u>AAGCTT</u>GTTCTCGTTGCTCGGCGTCAT | <u>ACTAGT</u>CCAGAAGAACGCGCACATTCA | 501 bp |
| Upstream *ΔMSMEG_2696* (MS111) | <u>GGTACC</u>CTGCGGTCGATGAGCGAGCT | <u>TCTAGA</u>CCCTGTGGCGAAGGTGAAGAA | 534 bp |
| Downstream *ΔMSMEG_2696* (MS110) | <u>AAGCTT</u>GGGCGCAGGCGTTCGACGAA | <u>ACTAGT</u>GGTCTGCTCGGCATGGTCGT | 500 bp |

The primers used to amplify DNA segments are listed. Nucleotide sequences recognized by the restriction enzymes used for cloning are underlined.
